# Geomagnetic field intensity as a cue for the regulation of insect migration

**DOI:** 10.1101/733782

**Authors:** Guijun Wan, Ruiying Liu, Chunxu Li, Jinglan He, Weidong Pan, Gregory A. Sword, Gao Hu, Fajun Chen

**Author notes:** Corresponding authors E-mail addresses (F.J. Chen).

## Abstract

Geomagnetic field (GMF) intensity can be used by some animals to determine their direction and position during migration. However, its role, if any, in mediating other migration-related phenotypes remains largely unknown. Here, we simulated variation in GMF intensity between two locations along the migration route of a nocturnal insect migrant, the brown planthopper *Nilaparvata lugens*, that varied by ∼5 μT (GMF_50μT_ vs. GMF_45μT_) in field intensity. After one generation of exposure, we tested for changes in key morphological, behavioural and physiological traits related to migratory performance including wing dimorphism, flight capacity and positive phototaxis. Our results showed that all three morphological and behavioural phenotypes responded to a small difference in magnetic field intensity between the simulated northern vs. southern locations in ways expected along the migratory route. Consistent magnetic responses in the expression of the phototaxis-related *Drosophila-like cryptochrome 1* (*Cry1*) gene and levels of two primary energy substrates used during flight, triglyceride and trehalose, were also found. Our findings indicate GMF intensity can be a cue that regulates the expression of phenotypes critical for insect migration and highlight the unique role of magnetoreception as a trait that can help migratory insects express potentially beneficial phenotypes in geographically variable environments.

## 1. Introduction

Organisms on Earth are perpetually surrounded by the geomagnetic field (GMF). Many animals have evolved the ability to sense or even utilize the GMF components of field intensity, direction, or both, although the details of the underlying biophysical mechanisms are still largely unknown [1, 2]. Two mechanisms, including magnetite-based magnetoreception and radical-pair-based magnetoreception, have been studied most in terrestrial animals [3-5]. In particular, evidence is accumulating for radical-pair-based magnetoreception, which is mainly based on a quantum process involving different spin states of radical pairs under the effects of magnetic interactions and spin-selective reactions, with cryptochromes (Crys) as a potential agent for radical-pair-based magnetoreception [3, 4, 6-8].

In addition to using GMF for orientation and navigation, animals can also exhibit other physiological and behavioural responses to changes in magnetic field intensity [9-15]. However, few studies have focused on responses of animals to changes in geomagnetic field intensity at levels that naturally vary in their environment [12, 16]. Most studies of the bioeffects of GMF on organisms in the last decade have been conducted by either shielding or enhancing the GMF, and have highlighted its importance in maintaining homeostasis [13, 15, 17]. Our work with migratory white-backed planthopper, *Sogatella furcifera*, showed that the absence of GMF can affect the expression of important migration-related behaviours, potentially through Cry-mediated hormone signaling. This includes changes in positive phototaxis which is critical for the ascent process, and flight capacity which is energetically-intensive and one of the main parameters used to assess migration ability [9, 18]. Additionally, previous work in migratory birds showed that changes in GMF information (both intensity and direction) over the migration route can affect their fuelling decisions and migratory restlessness [19-22]. As such, we speculated that changes in GMF intensity might play a more general underlying role in the regulation of insect migration.

Insects can conduct long-distance migrations in a short period of time [23, 24]. The nocturnal migratory brown planthopper, *Nilaparvata lugens*, has an annual long-distance migration in East Asia. Generally, their northward migration is initiated in spring from tropical and subtropical areas in Indochina to South China, followed by several subsequent waves of migration during the summer from South to North China, as well as to Japan and Korea Peninsula. Moreover, as they cannot overwinter in temperate East Asia, the insects then conduct a southward return migration through similar routes in the fall [25, 26]. Physiological responses to variation in the GMF have been demonstrated in *N. lugen*s [27]. Notably, both candidate magnetite crystals [28] and characterization of potential magnetic field targets, including Cry1, Cry2 [29], and iron-sulfur cluster assembly1 (IscA1) [30], have been systematically explored. Adult *N. lugens* also exhibit wing dimorphism associated with migration consisting of macropterous individuals with functional wings for seasonal migration and brachypterous individuals with vestigial wings and enhanced fecundity [31]. Given that outcomes and potential benefits of expressing specific migratory traits are predicted to change over the migratory route, we tested for changes in wing dimorphism, positive phototaxis, flight capacity, gene expression and flight physiology associated with variation in GMF intensity between the *N. lugens* emigration and immigration areas. We provide the first report of responses of migration-related traits to a magnetic field intensity change experienced by a long-distance migratory insect.

## 2. Material and methods

### Insect Stock

Adult *N. lugens* were collected from paddy fields at Nanjing, Jiangsu province of China during their south-to-north migration in the summer of 2018 and reared indoors on susceptible Taichung Native 1 rice seedlings at 25±1□, 70-80% RH and 14:10 h light: dark cycle (Dark during 1800-0800 hours) to establish a lab colony.

### Magnetic Fields and Insect Exposures

The GMF intensity generally ranges from ∼25μT at the magnetic equator to ∼65μT at the magnetic poles. We use two DC-type Helmholtz coil systems (External diameter: 1200 mm) to mimic the GMF intensity of two points on the migration route of *N. lugens*, Zhanjiang city (21°12’ 29” N, 110° 21’ 11” E; intensity: 45000 ± 255 nT, GMF_45μT_) and Nanjing city (32° 3’ 42” N, 118° 46’ 40” E; intensity: 50000 ± 239 nT, GMF_50μT_) at same declination (−5.7 ± 1.32°) and inclination (50.2 ± 1.23°). The GMF was altered in effective homogeneous areas of 300 × 300 × 300 mm^3^ inside each coil system. The parameters of the simulated magnetic field were monitored daily with a fluxgate magnetometer (Model 191A, HONOR TOP Magnetoelectric Technology Co., Ltd., Qingdao, China; sensitivity: ±1 nT). To ensure uniform environmental factors, the two coils systems were located in the same room separated by 6.5 m to avoid interference with each other. The position of tested *N. lugens* within the effective areas of the two coil systems was randomly changed in the same way daily. Following an established rearing protocol, rice planthoppers were exposed to the GMF_50μT_ vs. GMF_50μT_ treatments for one generation from mated F0 females to newly emerged F1 adults that were used in the assays [9]. We exposed *N. lugens* to the different GMF intensities for one generation because the insects were expected to undergo at least one more long-distance migration after F0 generation.

### Wing dimorphism and behavioural assays

Once the F1 adults emerged, individuals from each GMF group were identified to sex and wing form. Knowing that only macropterous adults can migrate, and most migrants are unmated [32], macropterous unmated adults from the GMF_50μT_ and GMF_45μT_ groups were respectively tested for positive phototaxis, and flight capacity. According to the responses of the closely-related migratory species *S. furcifera* to GMF compensation, and given that 2-day-old adult *N. lugens* begin to take off for migration [25], newly emerged to 3-day-old adults (D1-D3) and 2-day-old (D2) adults were investigated for positive phototaxis and flight capacity from 1800–0800 hours (subjective dark period), respectively. Phototaxis was assessed using a previously developed assay system for rice planthoppers [9]. To ensure that the only difference between the groups was the imposed magnetic field and considering that *N. lugens* would not all emerge on the same day, ten female or male adults of the same post-emergence age (D1, D2 or D3) were randomly selected from GMF_50μT_ or GMF_45μT_ treatments and assigned to each channel of the phototaxis testing apparatus under corresponding magnetic field conditions at 1745 hours (allowing 15 min for acclimation). The positive phototaxis testing procedure used was the same as in [9] except that insects that failed to move to the light the night before were not re-tested in this study. The final sample size for each D1-D3 test group was at least 90 (see sample sizes in the figure legends). To compare flight distances between the two GMF treatments, an infrared-beam based 8-channel flight mill system was constructed primarily with clear acrylic plexiglass to avoid any anomalies triggered by ferromagnetic materials (granted patent: ZL201620774681.1). The rice planthoppers could make a circular flight (diameter: 120 mm) while affixed to flight arm made from twisted copper wire (diameter: 2 mm) with the vertical axis for the attachment of the adult’s with as previously described [9]. Eight insects could be tested per night. An equal number of adults from each GMF treatment were randomly distributed to corresponding flight mills at 1745 hours (allowing 15 min for acclimation). Any insects that appeared unhealthy or escaped from their flight arms were excluded for further analyses. A total of fifteen 2-day-old adult *N. lugens* of each sex were tested in each GMF treatment group.

### Molecular and biochemical analysis

We selected the multifunctional *Crys* (*Cry1* and *Cry2*) genes in 1 to 3-day-old macropterous unmated adults for transcript expression analysis by quantitative real-time polymerase chain reaction (qRT-PCR) using *Arginine kinase* and *alpha 2-tubulin* as reference genes. Nine biologically independent pools containing ten adults were frozen at the same time of 1800 hours in liquid nitrogen (collected at 1745 hours and acclimated for 15 min) for each group divided by sex, magnetic field intensity and age, just before conducting the positive phototaxis experiment. Total RNA isolation, quality control, and concentration determination for total RNA, cDNA synthesis, and qRT-PCR were conducted as previously described [9]. Primer information can be found in the supplementary material (table S1). The content of primary flight fuels, including triglyceride (Jiancheng Bioengineering Institute, Nanjing, China) and trehalose (Suzhou Comin Biotechnology Co. Ltd., Suzhou, China) [33, 34], in nine biologically independent pools containing ten 2-day-old adults for each group frozen at the same time of 1800 hours in liquid nitrogen (collected at 1745 hours and acclimated for 15 min) just before the flight capacity test, was also determined using commercially available test kits.

### Statistical Analysis

All data were analyzed using SPSS 20 (IBM Inc., Armonk, NY, U.S.A.). Associations among categorical variables were assessed with the two-sided Cochran-Mantel-Haenszel chi-square test, while continuous variables were analyzed with general linear models (GLM). For continuous variables, Shapiro–Wilk test was used to test for normality (*P* > 0.05) and Levene’s test for the homogeneity of variances (*P* > 0.05), before an analysis of variance (ANOVA). In this study, all variables were separated by sex to investigate the main effect of GMF intensity. Therefore, we didn’t take sex as a fixed factor in GLM, and no post-hoc multiple comparison tests for the sampling time or test day were performed. We used a two-sided Cochran-Mantel-Haenszel test to investigate the association between GMF intensity (GMF_50μT_ or GMF_45μT_) and positive phototaxis stratified by test day (D1, D2, D3), and a two-way ANOVA to analyze the effects of the GMF intensity, sampling time (D1, D2, D3) and their interactions on gene expression levels of *Cry1* and *Cry2* (log(x + 1) transformation) at *α* = 0.05 for both females and males. When we found a significant association between the two categorical variables, a follow-up chi-square test (two-tailed) with Yates’s correction was performed to detail the association between GMF intensity and the positive phototaxis for females or males during the three test days. If a significant effect of GMF intensity or of interactions between the GMF intensity and sampling time was found on the gene expression levels, we used follow-up one-way ANOVA to compare the means for GMF_50μT_ and GMF_45μT_ at *α* = 0.05. A chi-square test (two-tailed) with Yates’s correction was performed to investigate the association between GMF intensity and wing dimorphism. A one-way ANOVA was used to test for the effect of GMF intensity on flight distance, as well as the content of primary flight fuels for female or male adults at *α* = 0.05, respectively. Effect sizes were estimated using Cohen’s *w* and partial *η*^2^ for chi-square test (small effect: *w* = 0.1; medium effect: *w* = 0.3; large effect: *w* = 0.5) and ANOVA (small effect: partial *η*^2^ = 0.01; medium effect: partial *η*^2^ = 0.06; large effect: partial *η*^2^ = 0.14), respectively, based on the benchmarks of Cohen (2013) [35].

## 3. Results and discussion

Exposure of *N. lugens* to variation in GMF intensities that mimicked those found at their emigration and immigration locations affected the frequency of wing dimorphism. Significantly more adult macropterous females (128.57%, *X*^2^_*1, 323*_ = 27.21, *P* < 0.001, *w* = 0.30) and males (51.23%, *X*^2^ _*1,367*_ = 17.24, *P* < 0.001, *w* = 0.22) developed under GMF_50μT_ vs. GMF_45μT_ (figure 1*a*.). The observed higher macropterous ratio, reflecting more migratory phenotype adults, would be expected in the northern habitats that must be colonized by annual immigration and experience higher GMF intensities relative to the overwintering site. This finding suggests that GMF intensity cues associated with latitude can be used by *N. lugens* to help balance the trade-off between flight capability and reproduction that is commonly observed in migratory animals [31]. For *N. lugens*, the relative importance of reproductive capacity over flight should be higher at lower latitudes with lower GMF intensity, because lower latitude areas serve as the overwintering site and source for the future migrations [25].

**Figure 1.**
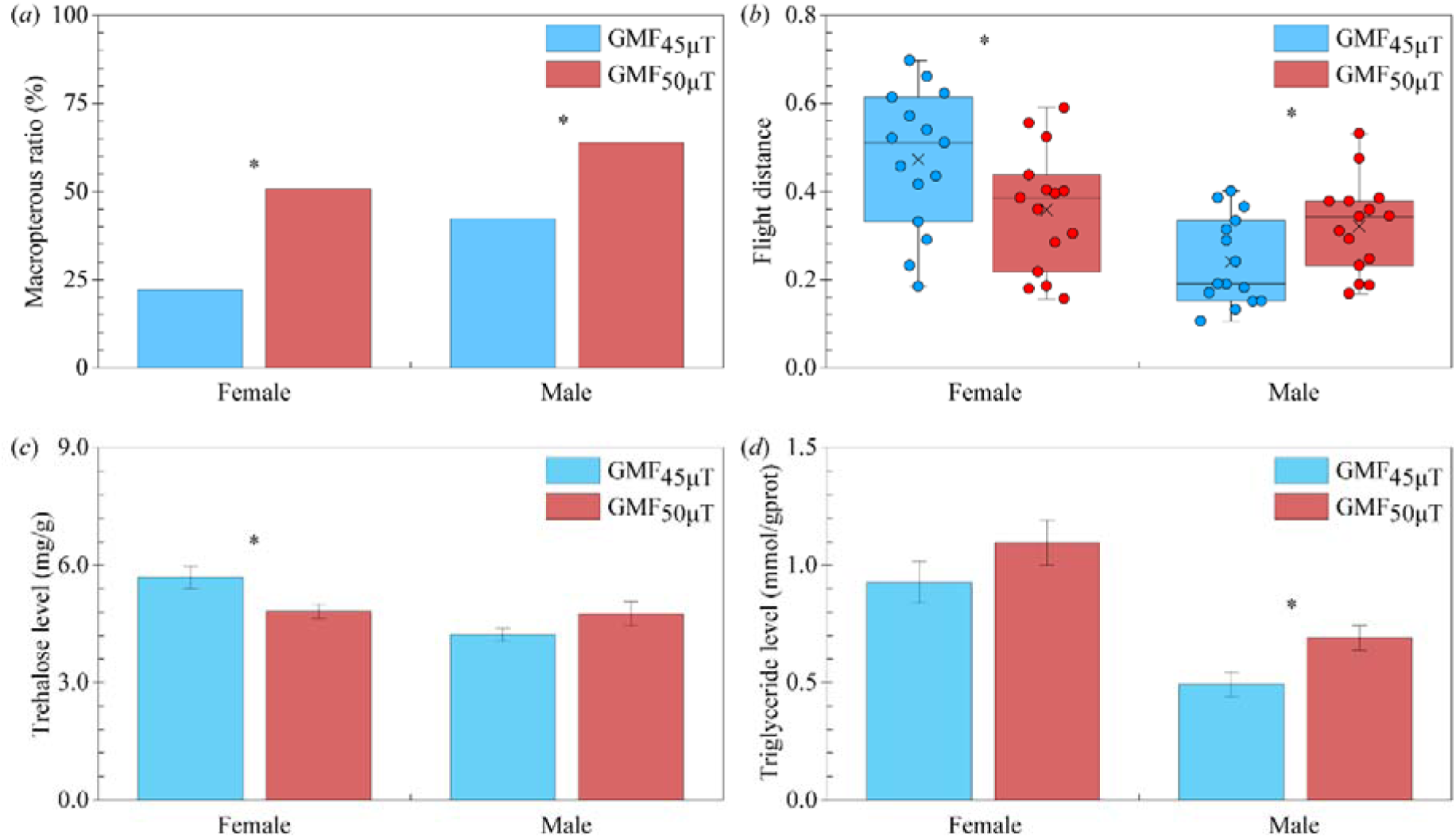
Wing dimorphism (*a*), flight distance (*b*), and main flight fuel trehalose (*c*) and triglyceride (*d*) levels of *Nilaparvata lugens* under GMF_50μT_ vs. GMF_45μT_. In (*a*), *N* = 156 and 167 for females, and 180 and 187 for males under GMF_50μT_ vs. GMF_45μT_, respectively. In (*b*), *N* = 15 for each group of unmated 2-day-old macropterous adults. In (*c*) and (*d*), samples of nine biologically independent pools containing ten unmated 2-day-old macropterous adults were used. * denotes significant association between GMF intensity and wing dimorphism by chi-square test with Yates’s correction (*a*), and significant differences between GMF_50μT_ vs. GMF_45μT_ by one-way ANOVA (*b*-*d*) for females or males at *p* < 0.05.

Interestingly, the simulated ∼5μT GMF intensity change potentially caused by long-distance migration also revealed a sexually dimorphic effect on flight capacity of 2-day-old adult *N. lugens*. Higher GMF_50μT_ vs. GMF_45μT_ significantly extended male flight distance (+33.84%; *F*_*1, 28*_ = 4.74, *P* = 0.038, partial *η*^2^ = 0.15) whereas it shortened that of females (−24.04%; *F*_*1, 28*_ = 4.45, *P* = 0.044, partial *η*^2^ = 0.14) (figure 1*b*). As one of the most important performance parameters associated with insect migration, our data suggest that male *N. lugens* could make longer flights than female adults during the south-to-north emigration. One possible reason for this sexual dimorphism in dispersal ability could be to relieve intraspecific competition by avoiding competition with their progeny and optimizing the use of resources. As main energy substances for insect flight, consistent significantly decreased trehalose level in females (−15.18%; *F*_*1, 16*_ = 5.89, *P* = 0.027, partial *η*^2^ = 0.27; figure 1*c*) and increased triglyceride level in males (+40.46%; *F*_*1, 16*_ = 6.33, *P* = 0.023, partial *η*^2^ = 0.28; figure 1*d*) were found under GMF_50μT_ vs. GMF_45μT_, indicating the potential pathway regulating trehalose and triglyceride (e.g., adipokinetic, insulin and juvenile hormone signaling) that is likely to be responsible for the sex-specific responses of flight distance to the GMF changes during migration[34, 36]. Taken together, the sexually dimorphic responses of flight capacity to the variation in the GMF likely reflects the different resource allocation strategies of females and males associated with long distance migration.

During the initial phase of migration, temporary inhibition of positive phototaxis was reported in *N. lugens* [37, 38]. In the first three days after eclosion that we monitored, unmated macropterous adults showed significantly weakened positive phototaxis under GMF_50μT_ vs. GMF_45μT_ (OR = 0.61, 95% CI: 0.50-0.76, *P* < 0.001, *w* = 0.11). Compared with GMF_45μT_, the higher GMF_50μT_ that would be experienced at northern sites significantly decreased the percentage of unmated macropterous females and males that moved towards the light (λ = 320-680 nm) by 30.39% (*X*^2^_*1, 660*_ = 7.05, *P* = 0.008, *w* = 0.11; figure 2*a*) and 19.49% (*X*^2^_*1, 880*_ = 7.92, *P* = 0.005, *w* = 0.10; figure 2*b*) during the first 3 days, respectively. Specifically, we found that GMF_50μT_ significantly decreased the move-to-light ratios of unmated macropterous females on the 2nd (−55.26%; *X*^2^_*1, 180*_ = 10.47, *P* = 0.001, *w* = 0.25) and 3rd day (−36.59%; *X*^2^_*1, 220*_ = 4.206, *P* = 0.040, *w* = 0.15; figure 2*a*), while males only on the 3rd day (−35.50%; *X*^2^_*1, 250*_ = 14.31, *P* < 0.001, *w* = 0.25; figure 2*b*). Although the biological significance of positive phototaxis is still an open question, it has been associated with migration in *N. lugens* [37,38] and the weakened positive phototaxis under a higher GMF intensity as would be expected in northern locations may help adult *N. lugens* smoothly ascend to the sky after take-off for their each wave of south-to-north migration without distraction by light sources on the ground [38].

**Figure 2.**
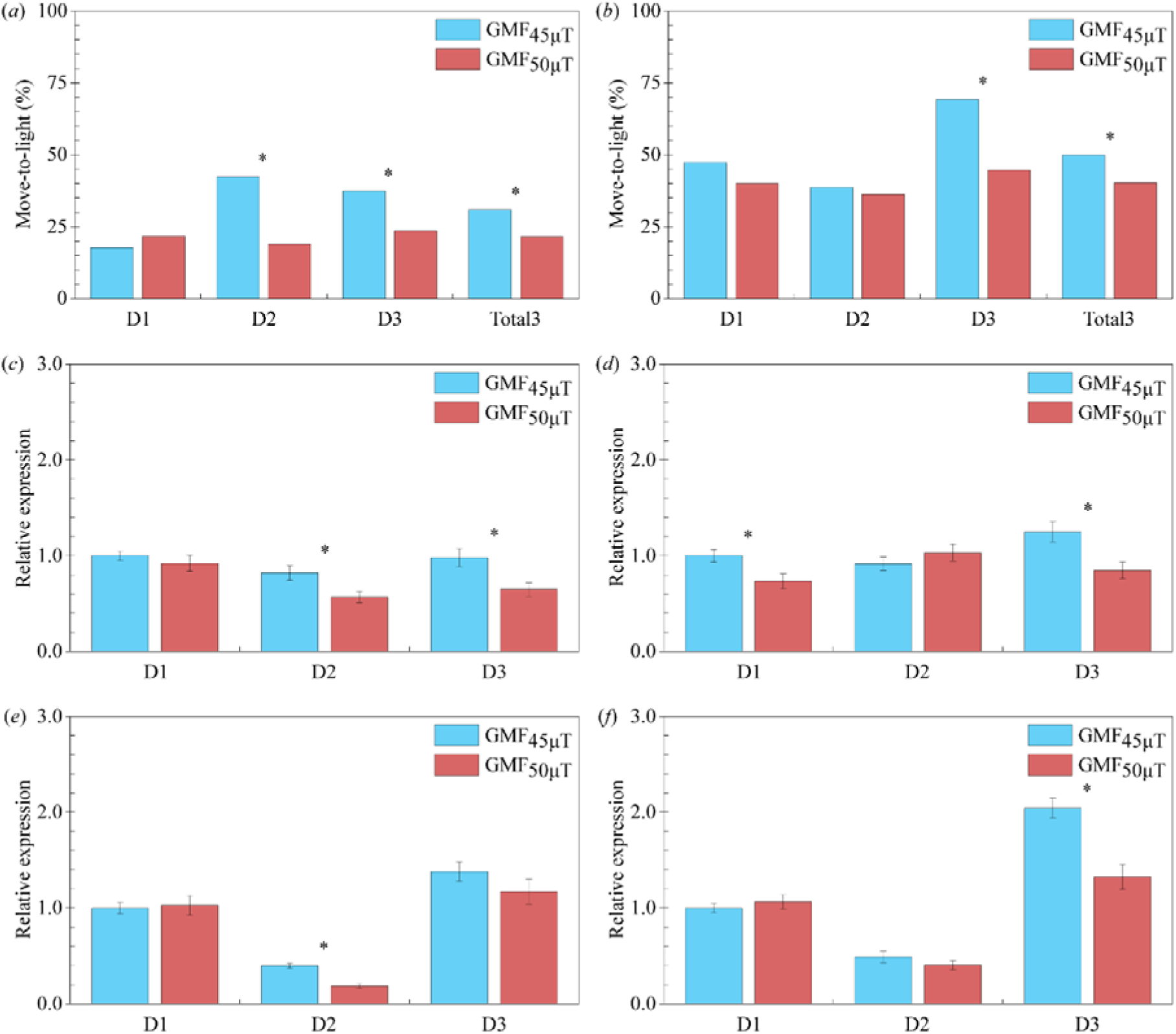
The positive phototaxis (*a, b*) and transcript expression analyses for the multifunctional *cryptochromes* (*Cry1* (*c, d*) and *Cry2* (*e, f*)) of unmated macropterous female (left panels) and male (right panels) *Nilaparvata lugens* under GMF_50μT_ vs. GMF_45μT_. For positive phototaxis, move-to-light percentage of adults on the 1st-3rd day (D1-D3) and in the first three days (Total3) after emergence were investigated. *N* = 130, 90, 110, 330, respectively, for the females in D1–D3 and Total3 under both GMF_50μT_ and GMF_45μT_; *N* = 130, 130, 130, 390, respectively, for the males in D1–D3 and Total3 under GMF_50μT_; *N* = 220, 150, 120, 490, respectively, for the females in D1–D3 and Total3 under GMF_45μT_. In (*c*) - (*f*), samples of nine biologically independent pools containing ten unmated macropterous adults were used. * denotes significant association between GMF intensity and positive phototaxis by chi-square test with Yates’s correction (*a, b*), and significant differences between GMF_50μT_ vs. GMF_45μT_ by one-way ANOVA (*c*-*f*) for each test (sampling) day at *p* < 0.05.

Extreme variation in GMF exposure has previously been shown to affect wing dimorphism, flight duration, and positive phototaxis in another rice planthopper species, *S. furcifera* [9]. However, the changes observed were induced by exposure to near-zero GMF. Importantly, our results here for *N. lugens* were induced by a biologically- and ecologically-relevant degree of variation in GMF levels between southern regions that serve as the source of migrants and northern regions where populations are re-established by migratory individuals every year. Thus, our results indicate that naturally-occurring variation in GMF intensity can affect the expression of traits involved in migration to result in phenotypes that favor different migratory and reproductive strategies in different locations as part of the annual migration cycle.

It is well established that multifunctional cryptochromes are involved magnetoreception, phototaxis, and the circadian clock [39, 40], and developmental changes in gene expression of adult *Cry1* and *Cry2* associated with extreme variation in the GMF have been previously observed in *N. lugens* [29]. In our study, except for *Cry1* in males, the expression levels of *Crys* in both adult females and males on the 1st-3rd days (D1–D3) after emergence all varied significantly over time (*F*_*2*_, μT 45μT _*48*_ ≥ 5.64, *P* ≤ 0.006, partial *η*^2^ ≥ 0.19). Significant interactions between the GMF intensity and sampling day on the expression levels of *Crys* were only found in males (*F*_*2, 48*_ ≥ 4.71, *P* ≤ 0.014, partial *η*^2^ ≥ 0.16), while the expression levels of *Crys* in both females and males were significantly affected by the GMF intensity (*F*_*1, 48*_ ≥ 6.14, *P* ≤ 0.017, partial *η*^2^ ≥ 0.11; Table 1). Mazzotta *et al.* (2013) found that the loss-of-function *Drosophila* Cry mutant can impair phototaxis of the fly [40]. Considering this, the consistent temporal expression pattern of *Cry1* may be responsible for the weakened positive phototaxis we observed. For both females and males, significantly decreased gene expression levels were observed in 2-(−30.78%; *F*_*1, 16*_ = 6.10, *P* = 0.025, partial *η*^2^ = 0.28) and 3-day-old (−34.09%; *F*_*1, 16*_ = 7.00, *P* = 0.018, partial *η*^2^ = 0.30; figure 2*c*) females as well as 1-(−26.67%; *F*_*1, 16*_ = 6.19, *P* = 0.024, partial *η*^2^ = 0.28) and 3-day-old (−32.17%; *F*_*1, 16*_ = 7.76, *P* = 0.013, partial *η*^2^ = 0.33; figure 2*d*) males under GMF_50_ vs. GMF. Significantly down-regulated *Cry2* expression was also found in 2-day-old females (−52.86%; *F*_*1, 16*_ = 29.36, *P* <0.001, partial *η*^2^ = 0.65; figure 2*e*) and 3-day-old males (−35.24%; *F*_*1, 16*_ = 16.75, *P* = 0.001, partial *η*^2^ = 0.51; figure 2*f*) adults. Unlike *Drosophila*-like Cry1, Vertebrate-like Cry2 has been reported to be vestigial flavoproteins [41] and would be unlikely to function as a magnetosensor by itself. Considering that *Drosophila*-like Cry1 is known to be involved in magnetosensing, it is possible in *N. lugens* Cry1 functions by itself for magnetoreception [1, 3, 4]. The magnetic response of positive phototaxis we found here is likely to be a result of interactions between the roles of *Drosophila*-like Cry1 in phototaxis and magnetoreception [1, 3, 4, 40], even though Cry2 may also function in a complex with photoreceptors or downstream in magnetoreception signaling [42]. Moreover, given that Cry2 is a component of the core circadian clock and it is reported that circadian clock is sensitive to changes in magnetic field intensity [11, 43], a *Cry2* function as a potential timing mechanism for migration based on GMF intensity might also exist [44].

**Table 1.** Two-way ANOVAs with the geomagnetic field (GMF) as the main factor and sampling day as the sub-factor on the gene expression levels of *cryptochromes* (*Cry1* and *Cry2*) for the unmated macropterous female and male adults.

| Sex | Genes | df | GMF<sup>a</sup> | df | Sampling day<sup>b</sup> | df<sup>c</sup> | GMF*Sampling day |
| --- | --- | --- | --- | --- | --- | --- | --- |
| Female | *Cry1* | 1,48 | 11.77/0.001 (0.20) | 2,48 | 5.64/0.006 (0.19) | 2,48 | 1.37/0.264 (0.05) |
|  | *Cry2* | 1,48 | 6.14/0.017 (0.11) | 2,48 | 98.15/<0.001 (0.80) | 2,48 | 2.09/0.135 (0.08) |
| Male | *Cry1* | 1,48 | 6.59/0.013 (0.12) | 2,48 | 2.21/0.121 (0.08) | 2,48 | 4.71/0.014 (0.16) |
|  | *Cry2* | 1,48 | 9.24/0.004 (0.16) | 2,48 | 108.50/<0.001 (0.82) | 2,48 | 7.35/0.002 (0.23) |

Results are given as *F* value/*P* value (partial $\eta^2$ ).
<sup>a</sup> GMF - the GMF<sub>50μT</sub> vs. GMF<sub>45μT</sub>.
<sup>b</sup> Sampling day - 18:00 on the 1st (D1), 2nd (D2) and 3rd (D3) day after the adult emergence.
<sup>c</sup> Ten individuals were randomly mixed as one sample.

## 4. Conclusion

Taken together, our results provide the first evidence that the expression of crucial migration-related traits such as wing dimorphism, flight capacity and phototaxis can respond to changes in local GMF intensity during long-distance migration, indicating that the GMF can act as a cue for the regulation of migratory phenotypes. *N. lugens* individuals seasonally migrate from their permanent southern overwintering regions into higher latitude northern regions where overwinter survival is not possible. Thus, behavioural, morphological and physiological changes that favor the development of more mobile migratory phenotypes are predicted to be favored in northern latitudes to increase the likelihood of successful return to the overwintering sites at the end of the season. Our results indicate that *N. lugens* phenotypes can vary in response to changes in GMF in a manner consistent with the expected trade-offs between reproduction and migration in permanent overwintering habitats in the south and ephemeral habitats in the north colonized annually as part of the seasonal migration cycle.

## Acknowledgments

We thank Likun Li for maintaining the flight mill and coils system, Jingyu Zhao and Jingjing Xu for their useful discussion.

