## Supplementary material for "Geomagnetic field intensity as a cue for the regulation of insect migration": Table S1

**Table S1.** Primers used to measure the transcript expression of *Cryptochrome1* and *Cryptochrome2* of unmated macropterous adult *Nilaparvata lugens* in the qRT-PCR assays.

| **Primer** | **Sequence (5’–3’)** | **Description** | **Genebank** |
| --- | --- | --- | --- |
| *NLAK*-F | TGCACATCAAAGTCCCCAAG | *Arginine kinase* (Reference gene) | KU365925.1 |
| *NLAK*-R | TCGTAGACACCGCCCTCG |  |  |
| *NLTub*-F | TGACCGAGTTCCAGACTAACCT | *Alpha 2-tubulin* (Reference gene) | FJ810204.1 |
| *NLTub*-R | AGACAACTGCTCGTGGTAGG |  |  |
| *NLCry1*-F | CAGACATGGGCTTCGATTTCA | *Cryptochrome 1* | KM108579.1 |
| *NLCry1*-R | ACCAGCACTTTCTCCGTCAAAT |  |  |
| *NLCry2*-F | CGCATACTCTCTACAGACTTGAT | *Cryptochrome 2* | KM108578.1 |
| *NLCry2*-R | CACCGTCTGGAATTTGCGATAC |  |  |
